## SUPPORTING INFORMATION for "COMBINED EFFECTS OF AMBIENT TEMPERATURE AND FOOD AVAILABILITY ON INDUCED INNATE IMMUNE RESPONSE OF A FRUIT-EATING BAT (*CAROLLIA PERSPICILLATA*)"

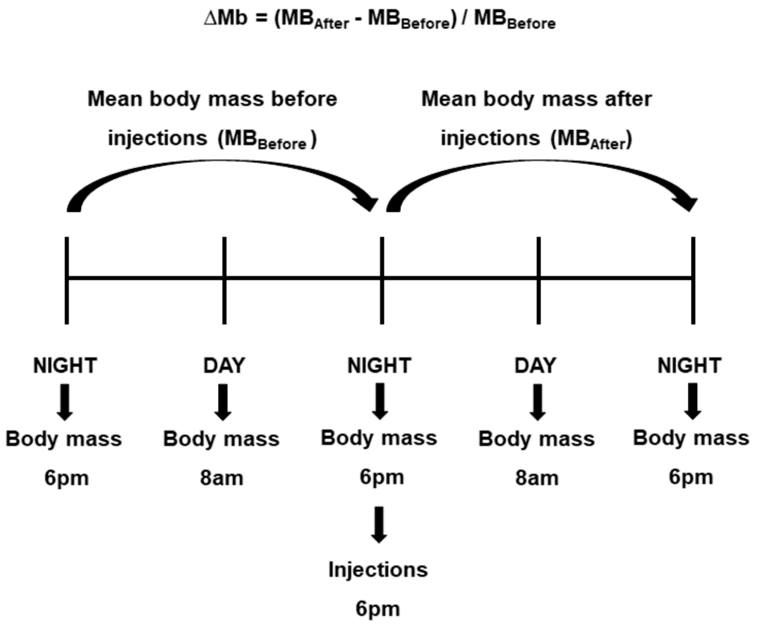


**S1 Fig.** Body mass change (ΔMb) of *C. perspicillata*. Body mass changes was assessed in relative terms as: body mass change (ΔMb) = (mean body mass after injections – mean body mass before injections) / (mean body mass before injections)

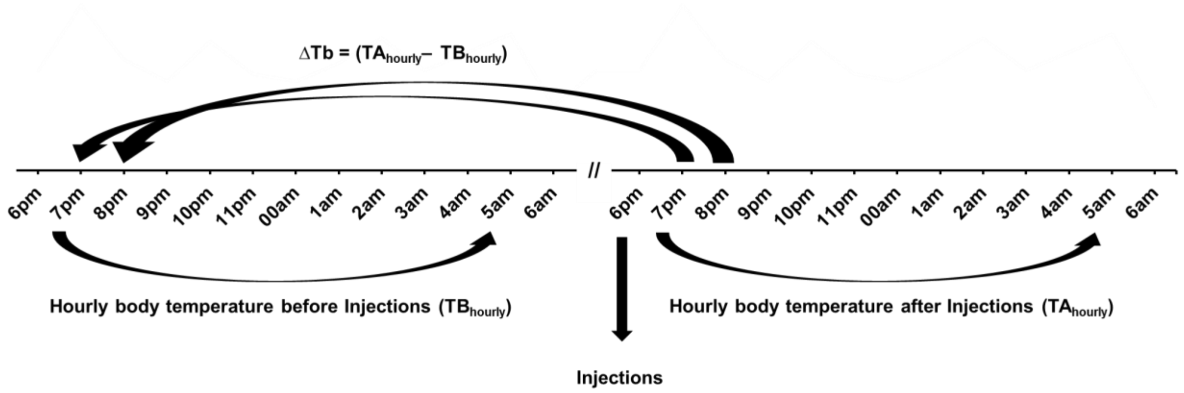


**S2 Fig.** Hourly body temperature change (ΔTb) of C. perspicillata. Hourly body temperature change was assessed 11 h after injections in absolute terms by subtracting hourly skin temperature after injections (TAhourly) from the respective hourly skin temperature before injections (TBhourly)

**S1 Table** Results of factorial ANOVA analyses to examine the effect of dose (PBS and 10 mg/kg LPS), ambient temperature (27°C and 33°C), diet (ad libitum feeding and restricted feeding) and the effects of their interactions on food intake change (ΔFI), body mass change (ΔMb), bacterial killing ability change (ΔBKA), total white blood cell change (ΔWBC) and neutrophil/lymphocyte ratio change (ΔNL).

| Variables | Factors | gl | F | Sig. | $\eta^2$ |
|---|---|---|---|---|---|
| Food intake Change (ΔFI) | Dose | 1 | 73.253 | **<0.001** | 0.567 |
| | Feeding Regime | 1 | 20.793 | **<0.001** | 0.271 |
| | Ambient Temperature | 1 | 2.980 | 0.090 | 0.051 |
| | Dose*Feeding Regime | 1 | 0.206 | 0.652 | 0.004 |
| | Dose*Ambient Temperature | 1 | 1.229 | 0.272 | 0.021 |
| | Feeding Regime*Ambient Temperature | 1 | 0.045 | 0.832 | 0.001 |
| | Dose*Feeding Regime*Ambient Temperature | 1 | 1.728 | 0.194 | 0.03 |
| | Error | 56 | | | |
| Body mass Change (ΔMb) | Dose | 1 | 56.713 | **<0.001** | 0.503 |
| | Feeding Regime | 1 | 25.31 | **<0.001** | 0.311 |
| | Ambient Temperature | 1 | 0.426 | 0.517 | 0.008 |
| | Dose*Feeding Regime | 1 | 0.001 | 0.98 | 0,000 |
| | Dose*Ambient Temperature | 1 | 0.134 | 0.716 | 0.002 |
| | Feeding Regime*Ambient Temperature | 1 | 0.069 | 0.794 | 0.001 |
| | Dose*Feeding Regime*Ambient Temperature | 1 | 0.025 | 0.874 | 0,000 |
| | Error | 56 | | | |
| BKA Change (ΔBKA) | Dose | 1 | 0.206 | 0.653 | 0.005 |
| | Feeding Regime | 1 | 0.568 | 0.455 | 0.014 |
| | Ambient Temperature | 1 | 0.28 | 0.599 | 0.007 |
| | Dose*Feeding Regime | 1 | 1.23 | 0.274 | 0.03 |
| | Dose*Ambient Temperature | 1 | 0.023 | 0.879 | 0.001 |
| | Feeding Regime*Ambient Temperature | 1 | 0.258 | 0.614 | 0.006 |
| | Dose*Feeding Regime*Ambient Temperature | 1 | 0.481 | 0.492 | 0.012 |
| | Error | 40 | | | |
| WBC Change (ΔWBC) | Dose | 1 | 1.375 | 0.246 | 0.024 |
| | Feeding Regime | 1 | 0.029 | 0.864 | 0.001 |
| | Ambient Temperature | 1 | 0.082 | 0.776 | 0.001 |
| | Dose*Feeding Regime | 1 | 0.020 | 0.887 | 0,000 |
| | Dose*Ambient Temperature | 1 | 0.688 | 0.41 | 0.012 |
| | Feeding Regime*Ambient Temperature | 1 | 0.209 | 0.649 | 0.004 |
| | Dose*Feeding Regime*Ambient Temperature | 1 | 0.295 | 0.589 | 0.005 |
| | Error | 56 | | | |
| N/L Change (ΔNL) | Dose | 1 | 157.691 | **<0.001** | 0.741 |
| | Feeding Regime | 1 | 7.789 | **0.007** | 0.124 |
| | Ambient Temperature | 1 | 4.792 | **0.033** | 0.08 |
| | Dose*Feeding Regime | 1 | 1.338 | 0.252 | 0.024 |
| | Dose*Ambient Temperature | 1 | 8.801 | **0.004** | 0.138 |
| | Feeding Regime*Ambient Temperature | 1 | 1.850 | 0.179 | 0.033 |
| | Dose*Feeding Regime*Ambient Temperature | 1 | 8.488 | **0.005** | 0.134 |
| | Error | 56 | | | |

**S2 Table** Food intake, body mass, total white blood cell count and neutrophil/lymphocyte ratio changes after immune challenge in *Carollia perspicillata*. **FIB**: mean food intake before injections; **FIA**: mean food intake after injections; **ΔFI**: mean food intake changes after injections; **WBCB**: mean total white blood cell count before injections; **WBCA**: mean white blood cell count after injections; **ΔWBC**: mean total white blood cell count change after injections; R**NLB**: mean neutrophil/lymphocyte ratio before injections; R**NLA**: mean neutrophil/lymphocyte ratio after injections; **ΔRNL**: mean neutrophil/lymphocyte ratio change after injections. **BMB**: mean body mass before injections; **BMA**: mean body mass after injections; **ΔMb**: mean body mass change after injections. N=7 for **ΔRNL** of food restricted group at 27°C injected with LPS. N= 8 for all other groups.

| Temperature | Feeding Regime | Treatments | FIB | FIA | ΔFIA (%) | BMB | BMA | ΔMb (%) | BKAB | BKAA | ΔBKA (%) | WBCB | WBCA | ΔWBC (%) | RNLB | RNLA | ΔRNL (%) |
|---|---|---|---|---|---|---|---|---|---|---|---|---|---|---|---|---|---|
| 27°C | Ad Libitum | PBS | 17.63 ± 2.93 | 16.66 ± 3.47 | -5.12 ± 13.69 | 13.87 ± 0.85 | 14.21 ± 1.09 | -1.22 ± 2.71 | 66.91 ± 6.74 | 65.99 ± 7.54 | -0.75 ± 13.49 | 4.14 ± 1.45 | 4.14 ± 1.15 | 2.18 ± 14.57 | 0.53 ± 0.13 | 0.54 ± 0.15 | 5.73 ± 35.94 |
| | | LPS | 15.28 ± 2.46 | 4.41 ± 2.62 | -69.89 ± 18.47 | 14.68 ± 1.04 | 13.54 ± 1 | -6.81 ± 2.38 | 70.92 ± 6.44 | 75.5 ± 5.98 | 5.41 ± 8.15 | 4.26 ± 1.26 | 5.17 ± 2.2 | 20.11 ± 36.4 | 0.55 ± 0.13 | 1.57 ± 0.3 | 200.44 ± 88.77 |
| | Resticted | PBS | 6.96 ± 0.69 | 12.38 ± 4.82 | 79.97 ± 74.09 | 14.38 ± 1.25 | 14.16 ± 0.93 | 2.31 ± 2.96 | 64.04 ± 13.62 | 72.38 ± 10.86 | 9.65 ± 17.65 | 4.14 ± 1.45 | 3.78 ± 1.25 | 4.79 ± 19.93 | 0.58 ± 0.13 | 0.58 ± 0.12 | 3.33 ± 24.3 |
| | | LPS | 7.26 ± 0.41 | 4.4 ± 3.96 | -39.7 ± 53.41 | 14.53 ± 0.72 | 14.23 ± 0.83 | -3.02 ± 4.36 | 65.24 ± 20.62 | 69.42 ± 10.99 | 10.63 ± 28.58 | 3.76 ± 1.7 | 4.03 ± 1.33 | 11.57 ± 17.94 | 0.74 ± 0.25 | 1.37 ± 0.41 | 57 ± 33.10 |
| 33°C | Ad Libitum | PBS | 10.76 ± 5.01 | 10.36 ± 4.48 | 12.86 ± 62.6 | 14.85 ± 1.33 | 15.04 ± 1.32 | -1.85 ± 2.37 | 70.34 ± 12.11 | 61.18 ± 17.44 | -4.09 ± 28.73 | 4.17 ± 1.8 | 4.65 ± 2.44 | 9.35 ± 23.41 | 0.48 ± 0.16 | 0.5 ± 0.07 | 11.12 ± 29.18 |
| | | LPS | 9.16 ± 3.5 | 0.63 ± 0.6 | -92.89 ± 5.94 | 14.63 ± 1.8 | 14.22 ± 1.73 | -6.71 ± 2.46 | 62.53 ± 15.59 | 66.04 ± 15.61 | 8.93 ± 26.11 | 3.72 ± 1.04 | 3.98 ± 1.1 | 8.53 ± 19.96 | 0.63 ± 0.14 | 1.98 ± 0.63 | 232.46 ± 150.34 |
| | Resticted | PBS | 7.33 ± 0.32 | 11.9 ± 4.29 | 64.21 ± 63.28 | 15.33 ± 0.96 | 15.08 ± 1.02 | 1.53 ± 1.63 | 55.5 ± 15.21 | 64.41 ± 12.13 | 8.43 ± 17.33 | 4.17 ± 1.8 | 3.81 ± 1.71 | 9.55 ± 14.07 | 0.61 ± 0.16 | 0.51 ± 0.12 | -10.2 ± 32.5 |
| | | LPS | 7.24 ± 0.42 | 3.31 ± 2.68 | -55.58 ± 34.87 | 15.24 ± 0.73 | 14.11 ± 0.73 | -3.51 ± 2.45 | 58.25 ± 13.28 | 60.36 ± 9.65 | -0.83 ± 22.49 | 3.86 ± 1.25 | 4.37 ± 1.32 | 16.59 ± 31.16 | 0.58 ± 0.22 | 1.75 ± 0.34 | 228.94 ± 107.48 |
| Mean between temperature and feeding regime treatments combined | | PBS | 10.67 ± 5.16 | 12.83 ± 4.72 | 37.98 ± 65.92 | 14.61 ± 1.2 | 14.62 ± 1.13 | 0.19 ± 2.95 | 63.67 ± 13.07 | 65.99 ± 12.23 | 3.45 ± 18.93 | 3.87 ± 1.5 | 4.09 ± 1.67 | 6.47 ± 17.78 | 0.55 ± 0.15 | 0.53 ± 0.12 | 2.5 ± 30.32 |
| | | LPS | 9.73 ± 3.92 | 3.19 ± 3.04 | -64.51 ± 37.4 | 14.77 ± 1.14 | 14.03 ± 1.12 | -5.01 ± 3.39 | 64.05 ± 14.93 | 68.14 ± 11.69 | 6.03 ± 21.69 | 3.9 ± 1.28 | 4.38 ± 1.55 | 14.2 ± 26.49 | 0.62 ± 0.2 | 1.67 ± 0.5 | 184.02 ± 122.54 |
| Mean between temperature and injections treatments combined | | Ad Libitum | 13.21 ± 4.86 | 8.02 ± 6.86 | -38.76 ± 54.75 | 14.87 ± 1.31 | 14.25 ± 1.36 | -4.15 ± 3.56 | 67.65 ± 10.6 | 67.45 ± 12.67 | 2.51 ± 19.47 | 4.07 ± 1.36 | 4.48 ± 1.8 | 10.04 ± 24.57 | 0.54 ± 0.15 | 1.15 ± 0.74 | 112.44 ± 136.64 |
| | | Resticted | 7.2 ± 0.48 | 8 ± 5.7 | 12.23 ± 82.62 | 14.51 ± 0.97 | 14.4 ± 0.93 | -0.67 ± 3.91 | 60.58 ± 15.61 | 66.64 ± 11.24 | 6.97 ± 21.03 | 3.7 ± 1.41 | 4 ± 1.36 | 10.63 ± 21.09 | 0.63 ± 0.2 | 1.05 ± 0.62 | 70.21 ± 113.69 |

Food intake changes (ΔFI) was assessed in relative terms as: food intake change = (food intake after injections – food intake before injections) / (food intake before injections); Body mass changes (ΔMb) was assessed in relative terms as: body mass change = (mean body mass after injections – mean body mass before injections) / (mean body mass before injections); Total withe blood cell changes (ΔWBC) was assessed in relative terms as: WBCs changes = (WBCs after injections – WBCs before injections)/ (WBCs before injections); Neutrophil/lymphocyte ratio changes (ΔNL) was assessed in relative terms as: N/L changes = (N/L after injections – N/L before injections)/ (N/L before injections); Bacterial killing ability (ΔBKA) assessed in relative terms as: BKA = (BKA after injections – BKA before injections)/ (BKA before injections)

**S3 Table** Results of factorial ANOVA analyses to examine the effect of dose (PBS and 10 mg/kg LPS), ambient temperature (27°C and 33°C), feeding regime (ad libitum and food restricted) and the effects of their interactions on body temperature change in *Carollia perspicillata*

| Variables | Factors | gl | F | Sig. | $\eta^2$ |
|---|---|---|---|---|---|
| Body temperature Change (ΔTb) | Time | 5 | 5.708 | **<0.001** | 0.093 |
| | Time*Feeding Regime | 5 | 1.490 | 0.197 | 0.025 |
| | Time*Ambient Temperature | 5 | 1.790 | 0.121 | 0.030 |
| | Time*Dose | 10 | 2.803 | **0.002** | 0.048 |
| | Time*Feeding Regime*Ambient Temperature | 5 | 1.607 | 0.164 | 0.027 |
| | Time*Feeding Regime*Dose | 5 | 1.093 | 0.363 | 0.019 |
| | Time*Ambient Temperature*Dose | 5 | 1.050 | 0.386 | 0.018 |
| | Time*Feeding Regime*Ambient Temperature*Dose | 5 | 0.637 | 0.658 | 0.011 |
| | Error | 258.33 | | | |
| | Feeding Regime | 1 | 0.213 | 0.645 | 0.003 |
| | Ambient Temperature | 1 | 1.739 | 0.192 | 0.030 |
| | Dose | 1 | 0.721 | 0.399 | 0.012 |
| | Feeding Regime*Ambient Temperature | 1 | 0.311 | 0.579 | 0.006 |
| | Feeding Regime*Dose | 1 | 3.018 | 0.087 | 0.051 |
| | Ambient Temperature*Dose | 1 | 0.451 | 0.504 | 0.007 |
| | Feeding Regime*Ambient Temperature*Dose | 1 | 0.992 | 0.323 | 0.017 |
| | Error | 56 | 0.001 | 0.985 | 0.000 |
| Time to reached the maximum increase in ΔTb | Feeding Regime | 1 | 4.079 | **0.050** | 0.127 |
| | Ambient Temperature | 1 | 1.751 | 0.196 | 0.058 |
| | Feeding Regime*Ambient Temperature | 1 | 0.819 | 0.373 | 0.028 |
| | Error | 28 | | | |

***** A factorial mixed ANOVA (3 Between subject factors / 1Within subject factor) was used to test effects of dose, ambient temperature, feeding Regime (as between subject factors), time (ΔTb for 11 hours as within subject factors) and the effects of their interactions on body temperature change (ΔTb). ** A factorial ANOVA was used to test effects of ambient temperature (27°C and 33°C) and feeding Regime (ad libitum and food restricted) and their interactions on the time at which the LPS-challenged groups reached the maximum increase in ΔTb.

**S4 Table** Hourly body temperature changes (ΔTb) changes after immune challenge in *Carollia perspicillata* kept at different ambient temperatures (27° and 33°C) and feeding regimes (ad libitum and food restricted). Data are given for all ambient temperature and feeding diets treatments combined.

| Temperature | Feeding Regime | Treatments | 1h | 2h | 3h | 4h | 5h | 6h | 7h | 8h | 9h | 10h | 11h |
|---|---|---|---|---|---|---|---|---|---|---|---|---|---|
| 27°C | Ad Libitum | PBS | -0.47±0.38 | -0.21±0.4 | -0.13±0.38 | -0.14±0.19 | -0.11±0.41 | -0.18±0.3 | -0.13±0.2 | -0.1±0.42 | -0.1±0.42 | -0.23±0.44 | -0.24±0.39 |
| | | LPS | -0.5±0.29 | -0.04±0.25 | 0.23±0.45 | 0.36±0.6 | 0.22±0.8 | 0.23±0.64 | 0.12±0.55 | 0.13±0.54 | -0.03±0.49 | -0.22±0.42 | -0.26±0.56 |
| | Resticted | PBS | -0.11±0.17 | -0.02±0.35 | -0.01±0.22 | 0.06±0.29 | 0.03±0.24 | 0.01±0.29 | 0.06±0.38 | 0.27±0.49 | 0.43±0.56 | 0.44±0.69 | 0.63±1.07 |
| | | LPS | -0.41±0.47 | -0.31±0.44 | 0.13±0.59 | 0.48±0.46 | 0.53±0.39 | 0.65±0.19 | 0.29±0.39 | 0.52±0.56 | 0.26±0.48 | 0.35±0.66 | 0.24±0.92 |
| 33°C | Ad Libitum | PBS | -0.38±0.52 | 0.07±0.36 | -0.11±0.62 | -0.16±0.19 | 0.2±0.27 | 0.21±0.25 | 0.07±0.51 | 0.04±0.41 | 0.04±0.26 | -0.03±0.28 | -0.07±0.57 |
| | | LPS | -0.06±0.73 | -0.17±0.57 | -0.03±0.99 | 0.36±0.78 | 0.38±0.65 | 0.12±0.56 | 0.02±0.72 | -0.1±0.49 | -0.16±0.75 | -0.02±0.65 | -0.2±0.6 |
| | Resticted | PBS | 0.07±0.65 | 0.04±0.46 | 0.13±0.57 | 0.03±0.3 | 0.04±0.29 | 0.09±0.24 | 0.03±0.26 | -0.04±0.64 | 0.08±0.34 | -0.11±0.66 | -0.18±1.12 |
| | | LPS | -0,14±0,43 | -0,43±0,47 | -0,29±0,29 | -0,11±0,49 | -0,01±0,87 | 0,12±0,55 | 0,22±0,58 | 0,19±0,61 | 0,04±0,89 | -0,25±1,22 | -0,34±1,2 |
| Mean between temperature and feeding regime treatments combined | | PBS | -0.22±0.49 | -0.03±0.39 | -0.03±0.47 | -0.05±0.26 | 0.04±0.32 | 0.03±0.29 | 0.01±0.35 | 0.05±0.5 | 0.11±0.44 | 0.02±0.58 | 0.04±0.88 |
| | | LPS | -0.28±0.52 | -0.24±0.45 | 0.01±0.63 | 0.27±0.61 | 0.28±0.7 | 0.28±0.53 | 0.16±0.55 | 0.18±0.57 | 0.03±0.66 | -0.04±0.79 | -0.14±0.85 |

Note Skin temperature change 11 h after injections was assessed in absolute terms (ΔTb) by subtracting hourly skin temperature after injections from the respective hourly skin temperature before injections

**S5 Table** Time at which the LPS-challenged groups reached the maximum increase in ΔTb

| Temperature | Feeding Regime | Time (h) |
|---|---|---|
| 27°C | Ad Libitum | 3.75±1.28 |
| | Resticted | 5.75±2.25 |
| 33°C | Ad Libitum | 6.38±2.97 |
| | Resticted | 6.75±3.20 |
| Mean between temperature treatments combined | Ad Libitum | 4.75±2.04 |
| | Resticted | 6.56±2.99 |
